# The immunological factors predisposing to severe COVID-19 are already present in healthy elderly and men

**DOI:** 10.1101/2021.04.30.442229

**Authors:** Gizem Kilic, Ozlem Bulut, Martin Jaeger, Rob ter Horst, Valerie A. C. M. Koeken, Simone Moorlag, Vera P. Mourits, Charlotte de Bree, Jorge Domínguez-Andrés, Leo A. B. Joosten, Mihai G. Netea

## Abstract

**Background:** Male sex and old age are risk factors for COVID-19 severity, but the underlying causes are unknown. A possible explanation for this might be the differences in immunological profiles in males and the elderly before the infection. Given the seasonal profile of COVID-19, the seasonal response against SARS-CoV-2 could also be different in these groups.

**Methods:** The abundance of circulating proteins and immune populations associated with severe COVID-19 was analyzed in 2 healthy cohorts. PBMCs of female, male, young, and old subjects in different seasons of the year were stimulated with heat-inactivated SARS-CoV-2.

**Result:** Several T cell subsets, which are known to be depleted in severe COVID-19 patients, were intrinsically less abundant in men and older individuals. Plasma proteins increasing with disease severity, including HGF, IL-8, and MCP-1, were more abundant in the elderly and males. The elderly produced significantly more IL-1RA and had a dysregulated IFNγ response with lower production in the summer compared with young individuals.

**Conclusions:** The immune characteristics of severe COVID-19, described by a differential abundance of immune cells and circulating inflammatory proteins, are intrinsically present in healthy men and the elderly. This might explain the susceptibility of men and the elderly to SARS-CoV-2 infection.

**Summary:** Immunological profile of severe COVID-19, characterized by altered immune cell populations and inflammatory plasma proteins is intrinsically present in healthy men and the elderly. Different age and sex groups show distinct seasonal responses to SARS-CoV-2.

## INTRODUCTION

Having emerged in China in December 2019, the coronavirus disease (COVID-19) caused by severe acute respiratory syndrome coronavirus-2 (SARS-CoV-2) has become a major health crisis. As of April 2021, SARS-CoV-2 has led to over 135 million infections and more than 3 million deaths worldwide [1].

The most vulnerable groups are people older than 70 years old and adults with underlying health conditions such as chronic respiratory problems and diabetes [2]. While age is the strongest predictor of death from COVID-19, the sharp increase in fatality after 50 years of age is more critical in men [3, 4]. Association of male sex with higher mortality has been consistently reported in different populations [5-7].

The factors underlying the impact of age and sex on the susceptibility to severe COVID-19 are, however, incompletely understood. Most studies to date have focused on the differences between either young and old or men and women during the disease process. A recent study has shown that male COVID-19 patients had higher circulating concentrations of cytokines such as IL-8 and IL-18, as well as a greater abundance of non-classical monocytes [8]. In contrast, a higher degree of T cell activation was observed in females than males during SARS-CoV-2 infection.

We previously analyzed 269 circulating proteins to identify COVID-19 severity markers by comparing patients in the intensive care unit (ICU) to patients who do not require ICU admission [9]. Several cytokines and chemokines such as IL-8 and monocyte chemoattractant protein-3 (MCP-3), and growth factors, e.g., hepatocyte growth factor (HGF), were increased in ICU patients, whereas stem cell factor (SCF) and several TNF-family proteins, e.g., TNF-related activation-induced cytokine (TRANCE) were decreased in ICU patients. Besides, frequently reported severe COVID-19 characteristics include elevated TNFα, IL-6, MCP-1, IP-10, and IL-10 concentrations, lower numbers and activity of T, B, and NK cells, lower antigen presentation, and downregulated type I interferon signaling [10-13]. Changes in cell populations and plasma proteins related to COVID-19 severity are also summarized in Supp. Table 1. However, it is unknown whether such differences are induced by the disease severity itself, or the potential to respond differently was already present in the healthy steady-state condition.

In this study, we analyzed some of the immune cell populations and circulating proteins linked to COVID-19 severity (Supp. Table 2) in two Dutch cohorts of healthy individuals. Our reasoning for choosing these parameters is based on the type of data available from the two cohorts. We investigated if healthy men and individuals over 50 years old already have an immunological profile that predisposes them to severe COVID-19 progression upon SARS-CoV-2 infection. Although the pandemic’s seasonal character is not completely clear, several studies reported links to temperature and humidity [14, 15]. Therefore, we also hypothesized that immunological differences induced by the seasons could influence the disease outcome. Hence, we investigated SARS-CoV-2-induced immune responses in healthy individuals *in vitro* at different time points of a year using cryo-preserved PBMCs from one of the cohorts. This study provides new insights into how age, sex, and seasons influence COVID-19 response (Figure 1).

**Figure 1.**
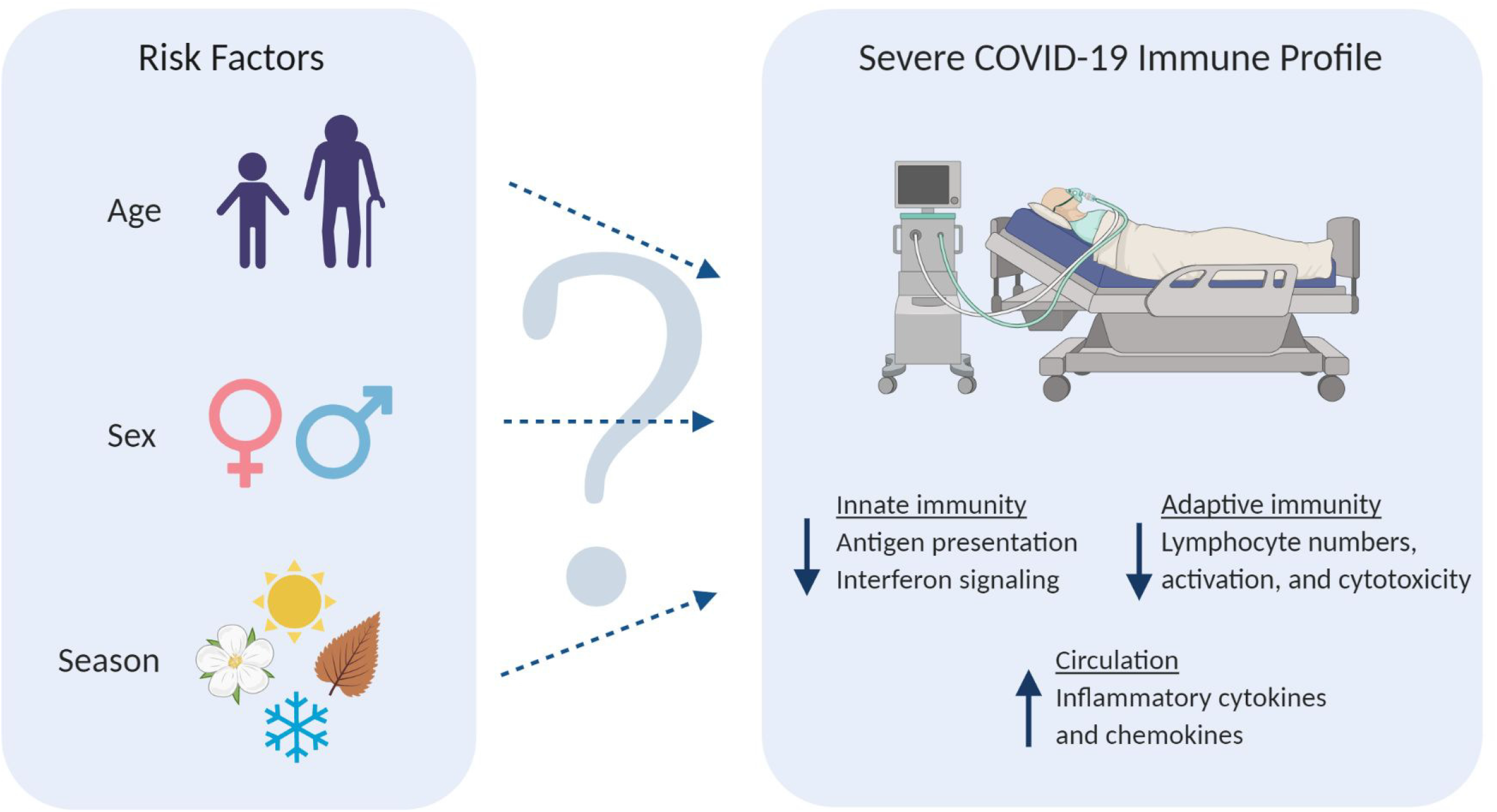
(Left) Potential COVID-19 risk factors investigated in this study. (Right) Frequently reported immunological characteristics of severe COVID-19 patients. Downward arrows depict a decline, while upward arrow represents an increase.

## MATERIALS & METHODS

### Study Cohorts

534 healthy individuals of Western European origin were included in Cohort 1 (500 Functional Genomics Project, see www.humanfunctionalgenomics.org) at the Radboud University Medical Center between August 2013 and December 2014. 45 volunteers were initially excluded due to medication use and chronic diseases, while 37 participants were later excluded from the analysis because one or more measurements were unavailable. Data of 452 participants, 229 females and 223 males with age ranges of 18-70 and 18-75, respectively, were used for analysis.

324 healthy individuals of Western European origin, Cohort 2, were included from April 2017 until June 2018 at the Radboud University Medical Center. 183 participants were female, and 141 were male with age ranges of 18-62 and 18-71, respectively. This cohort served as a validation cohort for proteomics.

Both studies were approved by the Arnhem-Nijmegen Medical Ethical Committee (NL42561.091.12 and NL58553.091.16). Inclusion and experimentation procedures were conducted according to the principles of the Declaration of Helsinki. Written informed consent was obtained from all volunteers before sample collection.

### Proteomics

Plasma proteins were measured using the Proximity Extension Assay (PEA) by Olink Proteomics (Uppsala, Sweden). The Olink Inflammation Panel consisting of 92 inflammation-related biomarkers was measured. This assay provides relative protein quantification expressed as normalized protein expression (NPX) values on a log2 scale. The proteins for which the missing data frequency was over 20 % were excluded from the analysis. The remaining data under the detection limit was replaced with the lower limit of detection for each protein. Measurements were normalized according to inter-plate controls.

### Flow Cytometry

Immune cell types in Cohort 1 were measured from whole blood by 10-color flow cytometry with Navios flow cytometer (Beckman Coulter, CA, USA). Staining and gating strategies were previously described in detail by Aguirre-Gamboa et al. [16].

### In vitro Stimulations and Cytokine Measurements

From the 452 individuals in Cohort 1, a sub-cohort of 50 people were asked to donate blood at 4 different time points in a year. Peripheral blood mononuclear cells (PBMCs) were collected and cryo-preserved between February 2016 and February 2017 to assess the seasonality of immune responses. From those 50, we selected 20 individuals considering the optimal age and sex matching of young-old and male-female comparisons. Therefore, cells isolated from 5 young males, 5 old males, 5 young females, and 5 old females were included for an *in vitro* study assessing seasonality’s impact on the cytokine responses to SARS-CoV-2. Cohort demographics are shown in Supp. Table 3. Upon thawing, PBMCs were stimulated with heat-inactivated SARS-CoV-2 at a concentration of 3.3×10^3^ TCID50/mL and heat-inactivated Influenza A H1N1 (California strain) at a concentration of 3.3×10^5^/mL for 24 hours and 5 days. Cytokine concentrations in the supernatants were measured with DuoSet^®^ ELISA kits (R&D Systems, MN, USA) according to the manufacturer’s protocols.

### Statistical Analyses

Statistical analyses were performed unsing R 3.6.1 (www.R-project.org) and GraphPad Prism 8 (GraphPad Software Inc., CA, USA). After adjusting the data for the covariate sex using linear regression, correlation of cell numbers or protein levels with age was done using Spearman’s rank-order correlation (Figures 2 and 4). Correction for sex was not applied in the heatmaps where two sexes were analyzed separately. After adjusting the data for the covariate age, differential protein expression or cell numbers between males and females was tested using the Mann-Whitney test (Figures 3 and 5). For box plots comparing different age groups, the two sexes, or four seasons, the Mann-Whitney test was used between any two groups. The Benjamini-Hochberg procedure was employed to correct multiple testing errors for the heatmaps and volcano plots. False discovery rate (FDR)-adjusted p-values smaller than 0.05 were considered statistically significant.

**Figure 2.**
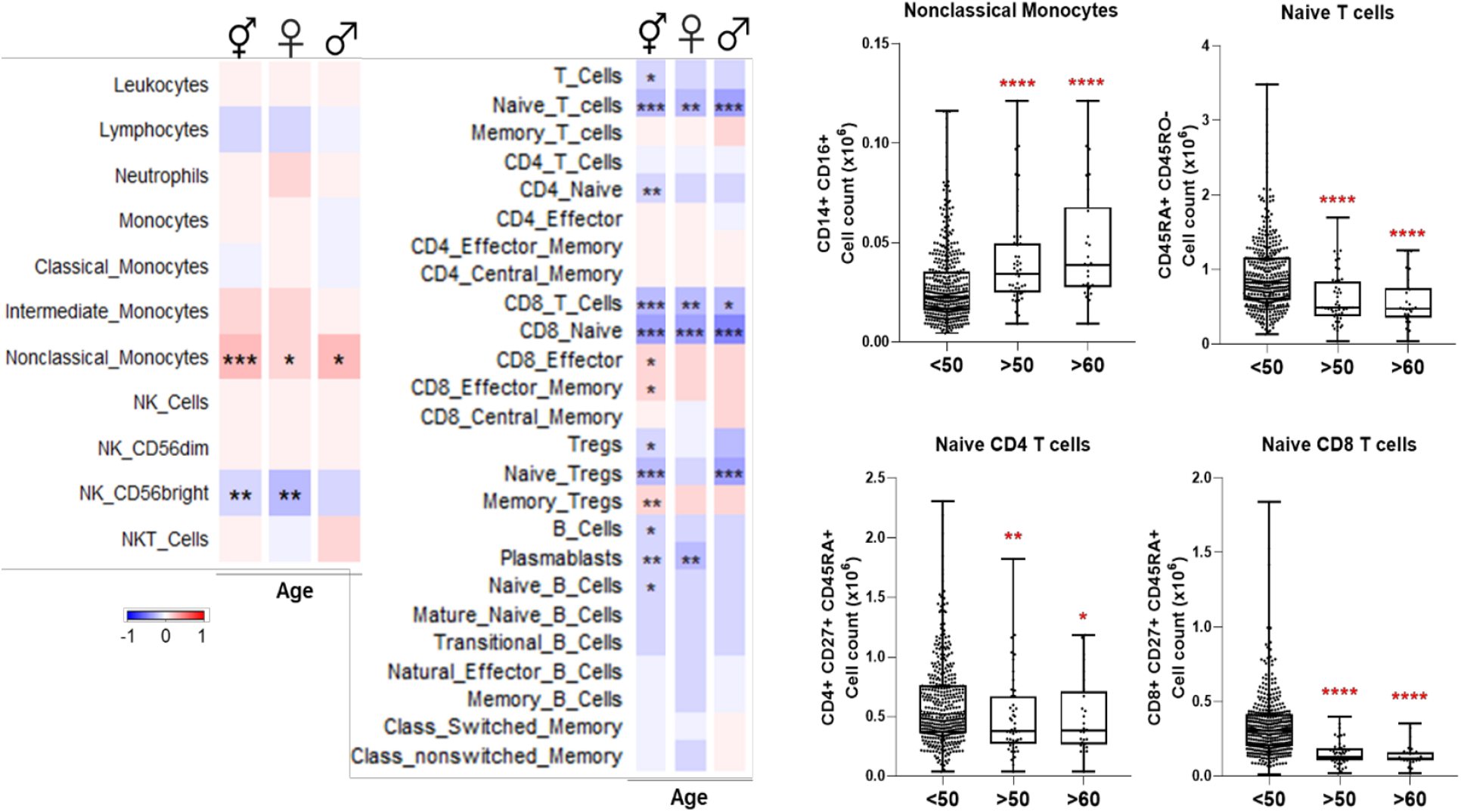
(Left) Heatmaps depicting Spearman correlation of age with cell numbers in Cohort 1. Red depicts positive correlation while blue depicts negative correlation. Data were controlled for the covariate sex when analyzing the whole cohort. (Right) Exemplary bar plots of selected cell populations in individuals aged less than 50, more than 50, and more than 60. *p ≤ 0.05, **p ≤ 0.01, ***p ≤ 0.001, ****p ≤ 0.0001. ⚥ whole cohort, ♀ females (n = 229), ♂ males (n = 223).

**Figure 3.**
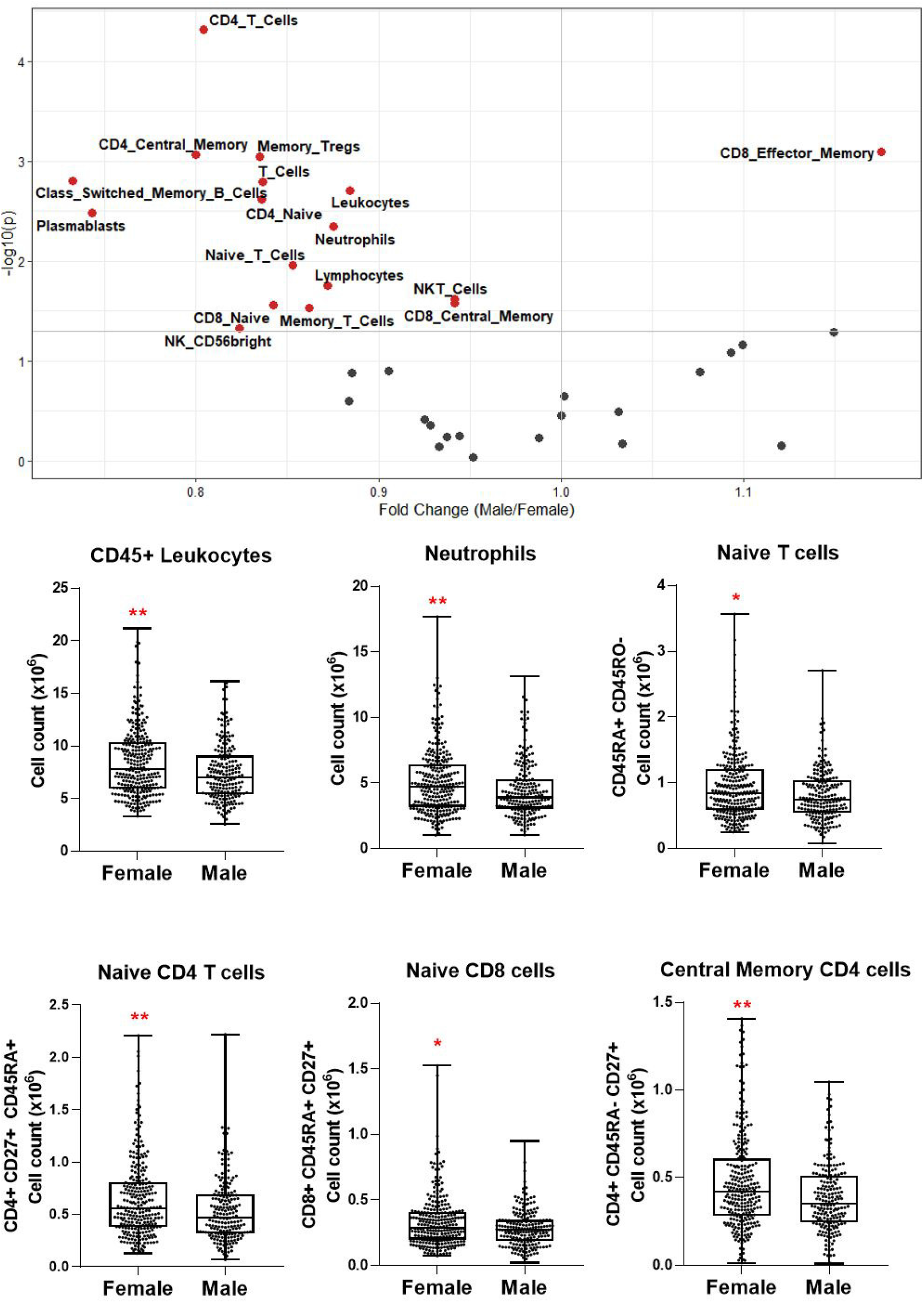
(Top) Volcano plot depicting differential numbers of cell populations depending on sex in Cohort 1. Significant results are depicted in red. Data were controlled for the covariate age. (Bottom) Exemplary bar plots of selected cell populations in females and males. *p ≤ 0.05, **p ≤ 0.01.

**Figure 4.**
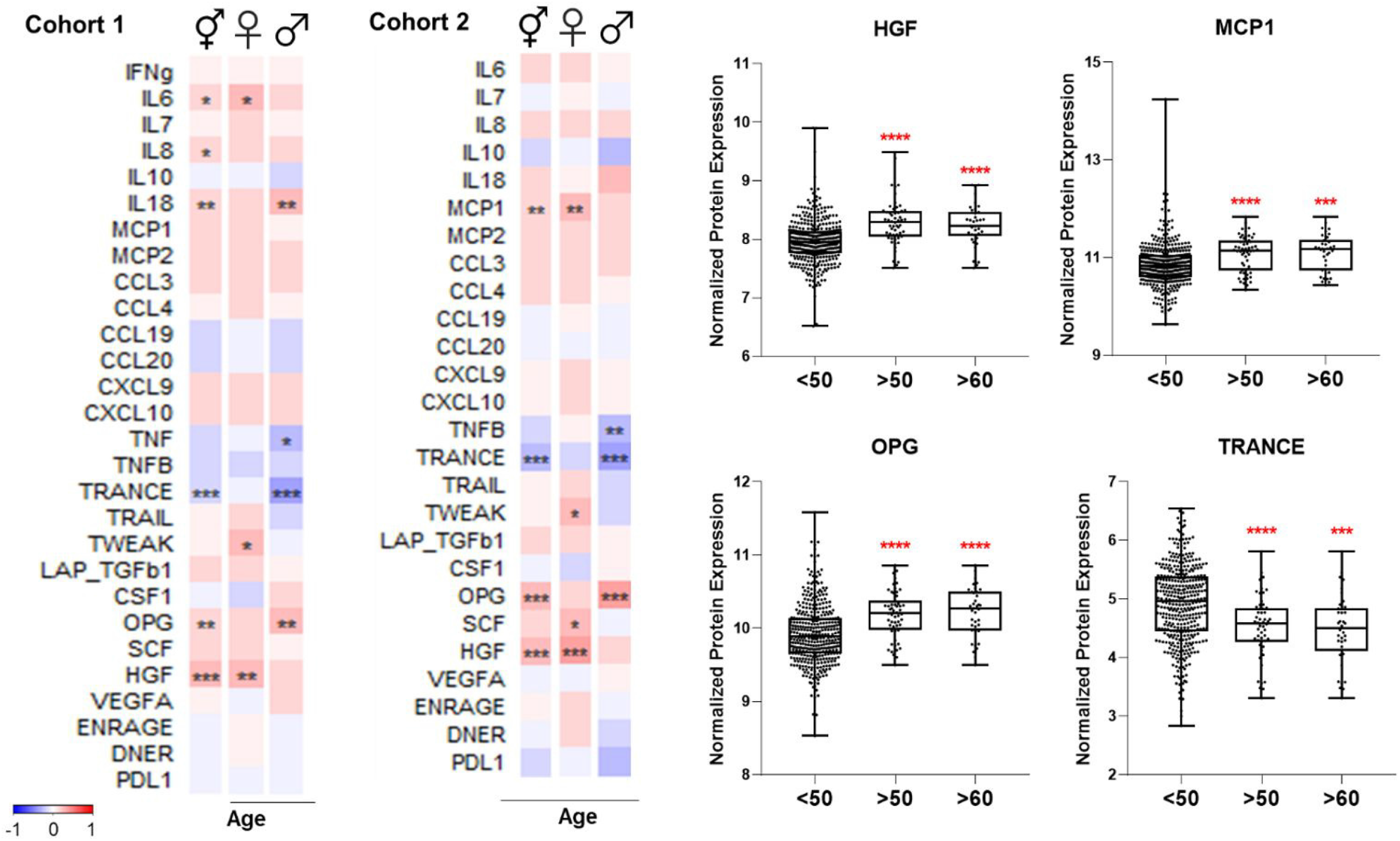
(Left) Heatmaps depicting Spearman correlation of age with plasma levels of proteins linked to COVID-19 severity. Red depicts positive correlation while blue depicts negative correlation. Data were controlled for the covariate sex when analyzing the whole cohort. (Right) Exemplary bar plots of plasma levels of selected proteins in individuals aged less than 50, more than 50, and more than 60. *p ≤ 0.05, **p ≤ 0.01, ***p ≤ 0.001, ****p ≤ 0.0001. ⚥ whole cohort, ♀ females (n_Cohort1_ = 229, n_Cohort2_ = 183), ♂ males (n_Cohort1_ = 223, n_Cohort2_ = 141).

**Figure 5.**
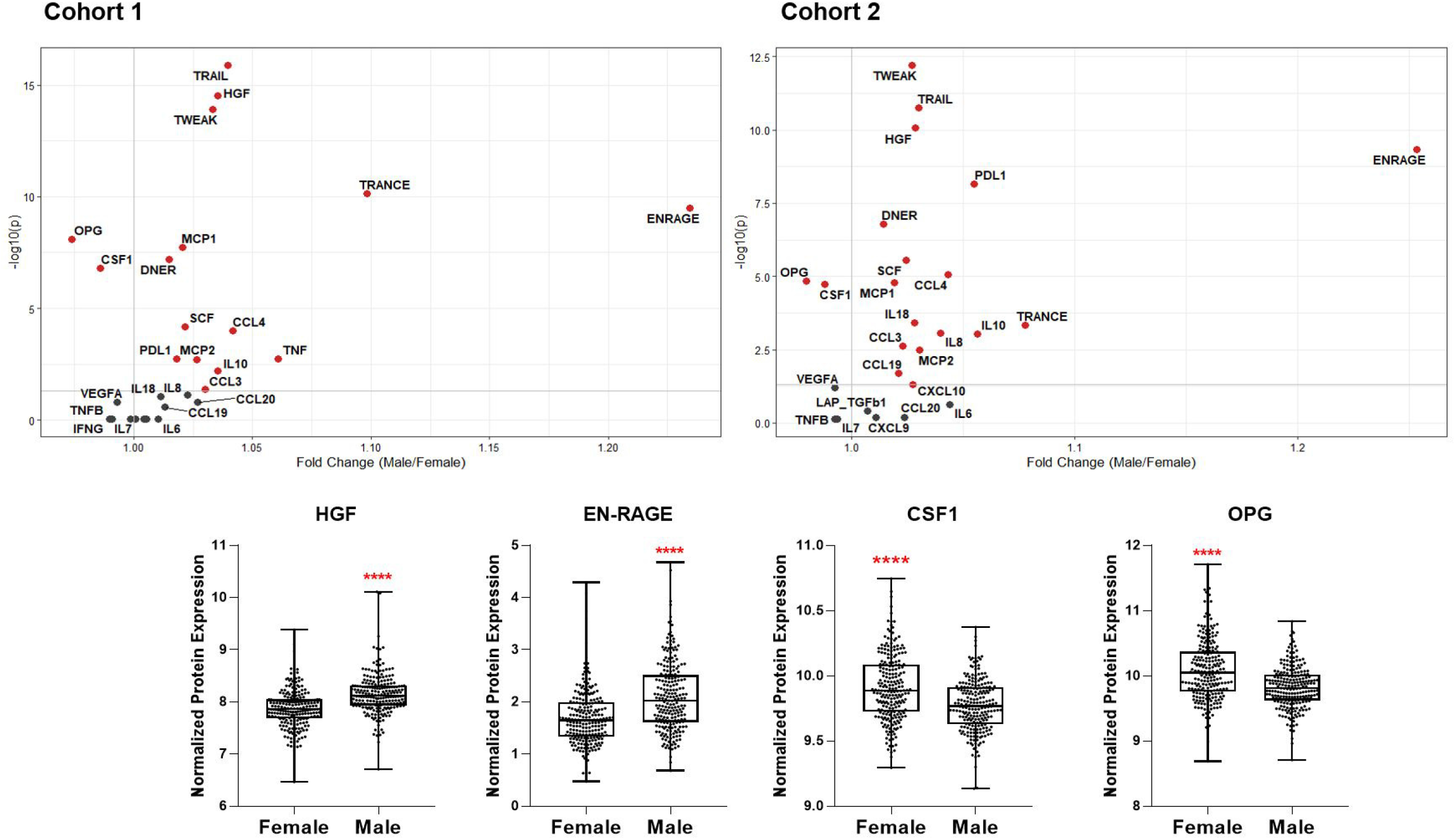
(Top) Volcano plots depicting differential plasma levels of proteins linked to COVID-19 severity depending on sex. Significant results are depicted in red. Data were controlled for the covariate age. (Bottom) Exemplary bar plots of selected proteins in females and males. ****p ≤ 0.0001.

## RESULTS

### Age-dependent Changes in Immune Cell Populations Linked to COVID-19 Severity

Various immune cell sub-types were correlated with age in Cohort 1. Non-classical monocytes increased with advancing age in both sexes, but intermediate and classical monocyte numbers were not significantly correlated with age. CD56^bright^ NK population decreased with age in females while other NK cell types remained unchanged (Figure 2). Previously, these results were partly reported by our group [16].

Total T cell count was negatively correlated with age, indicating the lymphopenia already experienced by the elderly, even in health, is likely to predispose to COVID-19 severity. Naïve T cells, especially CD8^+^, exhibited the most striking age-dependent decline in both sexes. Naïve regulatory T cells (Tregs) also decreased considerably with age in males, whereas memory Treg numbers were elevated. Total and naïve B cell numbers were negatively correlated with age. Plasmablasts similarly declined significantly in females. No significant age-related changes were observed among the other B cell sub-types.

Overall, these results demonstrate that naïve CD4^+^, CD8^+^, Treg, and B cell pools, as well as CD56^bright^ NK cells, which are all depleted in severe COVID-19, also decrease with age. The differences are clear even from the age of 50.

### Sex-dependent Patterns of Immune Cell Populations Linked to COVID-19 Severity

Next, we investigated the differential abundance of the same immune cell populations influencing COVID-19 severity in females vs. males. Only CD8^+^ effector memory T cells were significantly more abundant in males (Figure 3). Almost all cell types, including neutrophils, naïve CD4^+^ and CD8^+^ T cells, memory T cells, class-switched memory B cell, and CD56^bright^ NK cell counts, were significantly higher in females. The T cell types and CD56^bright^ NK cells, which are depleted in severe COVID-19 and elderly healthy people, are also apparently less abundant in males.

### Age-dependent Changes in Immune Mediator Proteins Linked to COVID-19 Severity

We selected 28 proteins whose plasma concentrations have been associated with severe COVID-19 and correlated them with age in healthy cohorts. Circulating IL-6 concentrations increased with age in healthy women, while IL-18 concentrations were higher in healthy men with advancing age in Cohort 1 (Figure 4). IL-8 concentrations positively correlated with old age in both females and males; however, it was only significant when sexes were combined. Among investigated chemokines, only MCP-1 was positively correlated with age in females in Cohort 2.

TNF-family proteins in plasma also changed with increasing age: TNF and TNFB concentrations were significantly lower in the circulation of older men in Cohort 1 and 2, respectively. TRANCE sharply declined in older individuals, more strikingly in males, while TWEAK was positively correlated only in females with advancing age. Moreover, aging in males was associated with elevated osteoprotegerin (OPG) concentrations, and HGF concentrations exhibited a considerable age-dependent increase in females in both cohorts.

In summary, several proteins in plasma that are increased in severe COVID-19 patients, such as IL-6, IL-8, IL-18, MCP-1, OPG, and HGF, are more abundant in healthy elderly compared to young individuals. Furthermore, proteins that are lower in severe COVID-19, e.g., TRANCE, decline with age.

### Sex-dependent Patterns of Immune Mediator Proteins Linked to COVID-19 Severity

We also compared plasma protein concentrations between sexes. It must be noted that Koeken et al. previously reported that males and females of Cohort 2 exhibit differences in baseline levels of many inflammatory markers [17]. Here we provide a more detailed analysis of COVID-19-related proteins among those in Cohorts 1 and 2. We observed a similar sex-dependent trend in both cohorts (Figure 5). Only OPG and colony-stimulating factor-1 (CSF-1), related to severe COVID-19, were significantly higher in women. On the other hand, plasma concentrations of other severity markers such as IL-8, IL-18, MCP-1, MCP-2, CCL3, and CCL4 were all higher in men. Furthermore, TRAIL, TWEAK, and TRANCE, which are all lower in COVID-19 patients in ICU, were more abundant in males [9]. Males exhibited more anti-inflammatory proteins, e.g., PD-L1 and IL-10. Growth factors HGF and SCF were also more abundant in male plasma.

These analyses show that most of the inflammatory mediators playing a role in infection severity are already higher in the circulation of healthy men.

We hypothesized that circulating inflammatory proteins could be related to impaired cytokine response against the virus. Therefore, we correlated SARS-CoV-2-induced *in vitro* cytokine productions with baseline circulating protein concentrations in a sub-cohort of Cohort 1. Indeed, PBMCs of the individuals with higher baseline plasma levels of MCP-2 and IL-8 produced more IL-1RA against SARS-CoV-2 *in vitro* (Supp. Figure 1). Furthermore, MCP-1 was negatively correlated with IFNγ production. The data indicate that higher baseline plasma concentrations of MCP-2, IL-8, and MCP-1 are associated with the inability to produce an optimal defense against SARS-CoV-2 infection.

### Sex, Age, and Season as Influencing Factors of Immune Response Against SARS-CoV-2

Next, we investigated the impact of seasonality on the SARS-CoV-2-induced immune response and assessed the contribution of age and sex. To this end, we selected 20 individuals from Cohort 1, for which cryo-preserved cells collected at 4 roughly equidistant time points in one year were available. We stimulated their PBMCs with heat-inactivated SARS-CoV-2 and influenza A H1N1.

We found that cytokine production upon SARS-CoV-2 stimulation did not substantially vary during the year, considering all 20 individuals (Supp. Figure 2). However, SARS-CoV-2-induced cytokine production did differ for different age groups and sexes throughout the year. Cytokines of the IL-1 biological pathway were higher in the elderly: IL-1β production tended to be greater in the elderly than in young individuals (Figure 6A), while SARS-CoV-2 induced more IL-1RA all-year-round in the old individuals(Figure 6B). Interestingly, IFNγ production upon stimulation with SARS-CoV-2 had a different seasonal profile in the young and elderly: young individuals produced more IFNγ in the summer and fall (Figure 6C). Remarkably, the elderly did not display this seasonal effect, with low IFNγ production throughout the year. TNFα and IL-6 productions upon stimulation were similar in the young and the elderly (Supp. Figure 3A-B). In addition, IL-1β production in response to SARS-CoV-2 in spring and summer was higher in females; however, the average yearly response failed to reach statistical significance (Figure 6D). IL-1RA and IFNγ production in males and females were comparable (Figure 6E-F).

**Figure 6.**
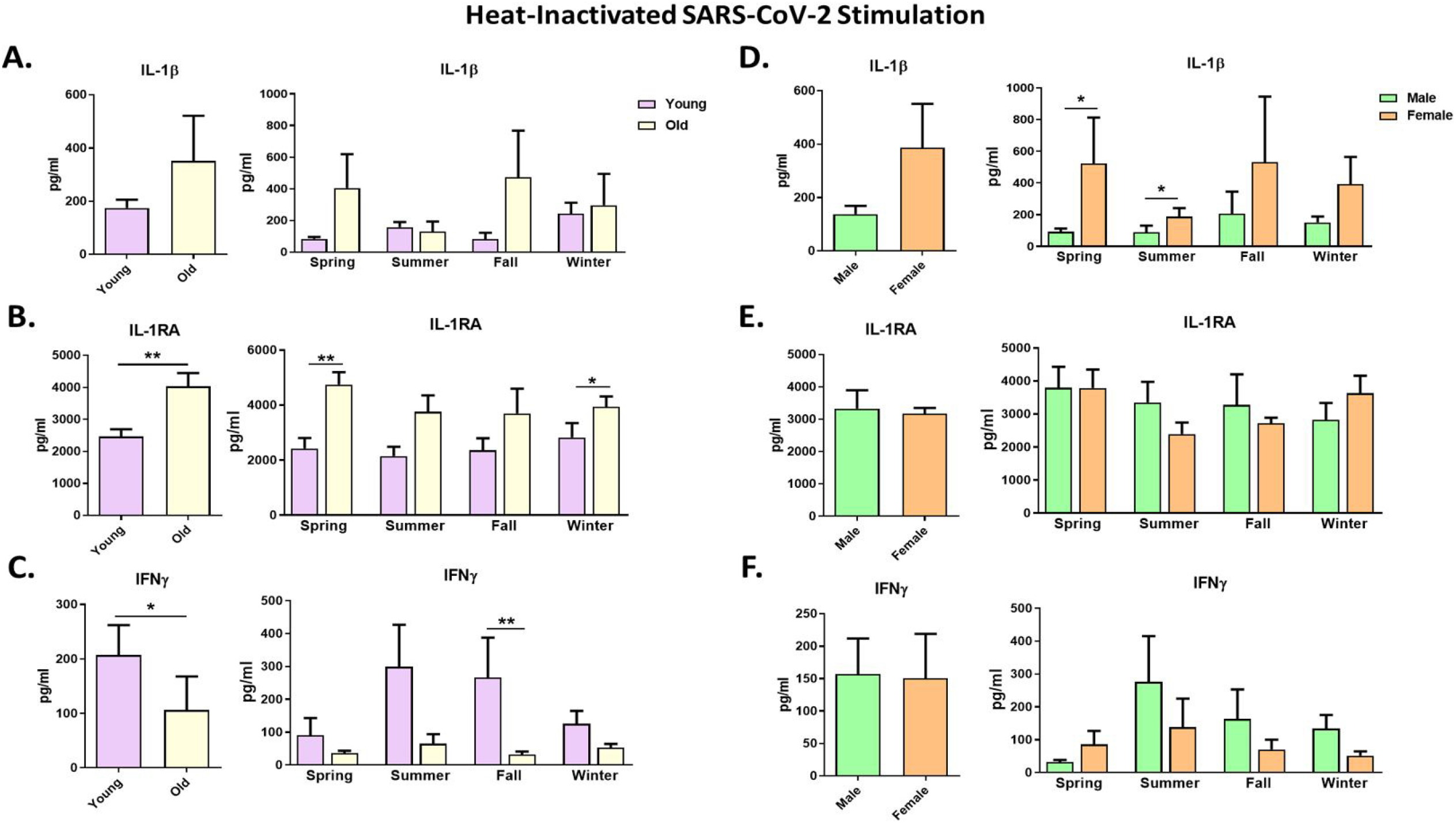
Cytokine responses against heat-inactivated SARS-CoV-2 in healthy individuals. Each panel’s left graphs show the yearly average production, while the right graphs demonstrate cytokine production in every season. Responses were compared between young and old individuals (A-C) and between males and males (D-F). IL-1β and IL-1RA cytokine levels were measured after 24 hours, while IFNγ was measured after 5 days. *p ≤ 0.05, **p ≤ 0.01. n=8-10. Error bars depict the standard error of the mean (SEM).

Stimulating PBMCs using another RNA virus, influenza H1N1, resulted in a similar pattern to SARS-CoV-2 stimulation regarding the age and sex effects. The elderly tended to produce more IL-1β and IL-1RA, while young individuals could produce higher IFNγ amounts upon influenza stimulation (Supp. Figure 4). Similar amounts of IL-1β, IL-1RA, and IFNγ were induced in males and females on average, with few exceptions (Supp. Figure 4D-F).

These data show that individuals of distinct ages and sexes respond differently to SARS-CoV-2 infection depending on the seasons of the year.

## DISCUSSION

Old age and male sex are important risk factors for COVID-19 severity. Several studies have investigated whether immune responses during SARS-CoV-2 infection are influenced by demographic factors such as age and sex [18, 19]. Although they identified immune profiles associated with severe COVID-19, they could not assess whether these were secondarily induced by the disease, or due to *a-priori* immune differences between different groups. In the present study, we show that the immune characteristics associated with severe COVID-19, such as specific changes in cell populations and circulating inflammatory proteins, are already present in healthy elderly and men. Interestingly, while the season did not impact the immune response to SARS-CoV-2 stimulation in the entire group, there was a clear difference in the responses between the young and old. Young individuals, but not the elderly, improve their IFNγ responses to SARS-CoV-2 during the summer.

COVID-19 progression to a severe clinical picture is related to the depletion of several immune cell types, including naïve CD4^+^ and CD8^+^, and CD56^high^ NK cells. Our study demonstrates the age-related decline of these cells in healthy individuals even before the infection, which likely contributes to their incapacity to eliminate the virus. These data are supported by studies suggesting that some of these cell types are scarcer in uninfected elderly and males [20, 21]. Naïve B lymphocyte numbers were also reduced with advanced age, which would undermine the development of adaptive immunity and antibody production upon infection [22]. The aging process does not only alter cell numbers, but also the functions (Figure 6C) [23]. All these might cumulatively disrupt the response against SARS-CoV-2 infection.

We showed that one of the striking differences in immune cell types between males and females was the number of CD4^+^ T cells. Although we cannot rule out that significant differences in cell populations might not determine the disease severity, SARS-CoV-2-specific CD4^+^ T cells were strongly linked with milder COVID-19, unlike antibodies and CD8^+^ T cell numbers [24]. Fast induction of CD4^+^ T cells was related to a milder disease, while defects in inducing SARS-CoV-2-specific CD4^+^ T cells were associated with severe or fatal COVID-19.

An exaggerated systemic inflammation has been associated with severe COVID-19, mirrored by high circulating concentrations of pro-inflammatory mediators [25]. Among those, IL-8, IL-18, and MCP-1 have been frequently reported, and the first two characterize the immune response of men with a severe outcome [8]. Notably, their concentrations are already higher in both men and the elderly in our healthy cohorts. Another protein displaying the same pattern is HGF, acting on epithelial and T cells promoting migration [26], which has higher concentrations in the circulation of severe patients [27]. The circulating HGF concentrations are higher in men, but also increases with age in women.

Chemokines are critical inflammatory mediators, and MCP-2, CCL3, CCL4, CCL19, and CXCL10 are all more abundant in males. ENRAGE (S100A12), produced by neutrophils and monocytes, is also higher in healthy males than females. Monocytes expressing high S100A12 and IL-8 are linked to COVID-19 severity [28]. Additionally, anti-inflammatory proteins IL-10 and PD-L1 are elevated in healthy males and severe COVID-19. Initially considered a negative feedback mechanism for infection-induced inflammation, there are arguments suggesting these proteins as biomarkers of immune exhaustion, which is likely to play a substantial role in the pathophysiology of COVID-19 [29, 30]. Notably, the association of early IL-10 production with COVID-19 severity supports this idea [31].

Severity markers increasing with old age, but not affected by sex, include IL-6 and OPG. IL-6 secreted by hyperactive monocytes contributes to low HLA-DR expression and lymphopenia in severe COVID-19 [32]. TNF-family cytokine receptor OPG, abundant in ICU patients, increases with old age in our healthy cohorts [9]. High OPG concentrations in females are likely due to estrogen’s effects promoting OPG expression to inhibit bone resorption [33]. Another TNF-family member, TRANCE (RANKL), which is lower in severe COVID-19 cases, also declines with advancing age in healthy individuals. T cells are one of the primary TRANCE sources, which might explain its scarcity in the elderly and severe COVID-19 patients with lymphopenia [34].

Of note, concentrations of circulating IL-7 and IFNγ, which are increased in severe COVID-19, are similar in men and women. Therefore, T cell numbers being higher in women is unlikely due to IL-7-induced lymphopoiesis, whereas higher T cell numbers do not necessarily lead to more circulating IFNγ. An overview of the age- and sex-dependent immune profiles in healthy individuals potentially predisposing to severe COVID-19 upon infection is provided in Figure 7.

**Figure 7.**
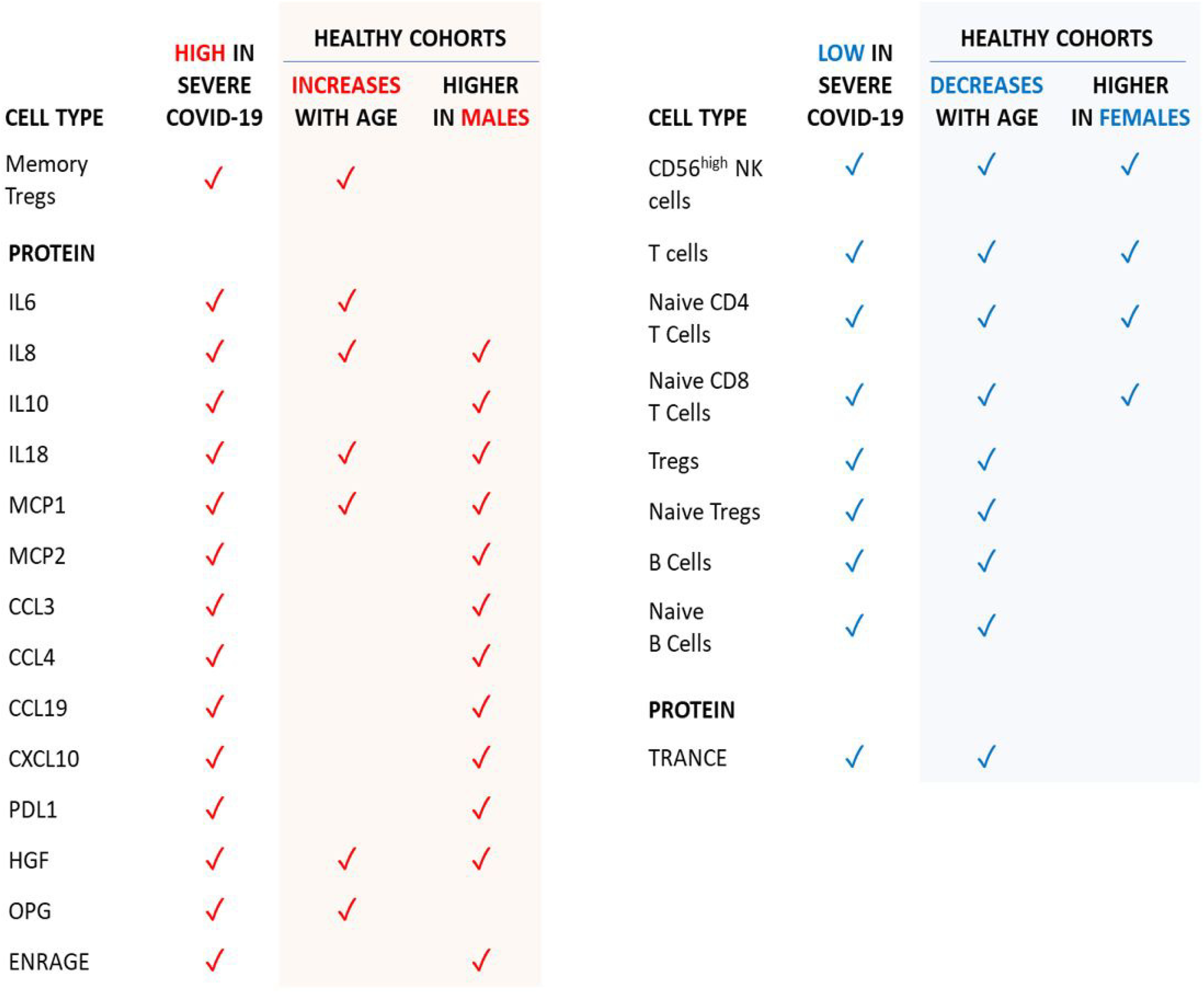
Age- and sex-dependent factors in healthy individuals that are in-line with the severe COVID-19 phenotype.

Environmental factors are also known to affect immune responses. Certain infections such as influenza follow a seasonal pattern [35], and the evolution of the pandemic last year also suggested that COVID-19 incidence might follow a seasonal variation [14]. Therefore, we questioned whether this might be due to seasonal changes in the immune response to the virus. In the entire group, we found no clear seasonal response against SARS-CoV-2, although this could be due to the limited sample size. Interestingly, young individuals improve their IFNγ response to SARS-CoV-2 during the summer, while the elderly do not. Further research with larger cohorts is required to validate seasonality’s full impact on anti-SARS-CoV-2 host defense.

Our results indicate that the immune response upon *in vitro* SARS-CoV-2 stimulation varies depending on age and sex. The response of the elderly is characterized by low IFNγ and elevated IL-1RA production. IFNγ is crucial for an effective response of T- and NK cells to viral infections, and its deficiency is associated with severe COVID-19 [36]. Depleted NK and T cell pools might explain why the elderly are less capable of producing IFNγ upon SARS-CoV-2 stimulation. Poor IFNγ response, especially in the fall, might put the elderly at higher risk for severe COVID-19. Men and women produce comparable amounts of IFNγ, although the T cell numbers are higher in women. Other roles of T cells besides IFNγ production may contribute to the better prognosis of women with COVID-19. One important point is that these defects are present in the whole population, not at the individual level: while the elderly as a group have lower immune responses, there are certainly aged individuals who have effective immune reaction. This inter-individual variability could be the reason why some elderly or some men have good responses against SARS-CoV-2 and develop only mild disease.

In COVID-19, an overproduction of pro-inflammatory cytokines contributes to the pathophysiology late in the disease [37]. On the other hand, cytokines such as IL-1β might also be crucial for an early anti-viral response. The deficiency of IL-1β or its receptor causes higher viral load and mortality in murine models [38]. Furthermore, genetic variants in *IL1B* contribute to influenza susceptibility in humans [39]. We observed higher IL-1β production in women in response to SARS-CoV-2 *in vitro*, arguing that an initial potent anti-viral defense is essential to prevent severe disease. Moreover, IL-1RA is an antagonist of IL-1 bioactivity, helping prevent excessive inflammation [40]. However, early IL-1RA production in patients is associated with COVID-19 severity [31], while our finding of higher IL-1RA production in the elderly suggests that IL-1RA might be hindering their ability to mount an optimal immune response against SARS-CoV-2. Alternatively, high IL-1RA production with increasing age might mirror the general inflammatory profile of elderly individuals.

In conclusion, our findings shed light on the immunological factors that might explain why men and the elderly have a higher risk of developing severe COVID-19. These results also emphasize the importance of the IL1β/IL-1RA axis and IFNγ in anti-SARS-CoV-2 response. We propose that intrinsically different immune characteristics, including plasma inflammatory mediators and immune cell populations, in healthy people would influence their immune response upon SARS-CoV-2 infection and the severity of the disease. Results of this study inform prophylactic and therapeutic efforts.

## Results were partly presented in

Second Biomarker Meeting of Paris on November 25^th^ 2020 (virtual).

## REFERENCES

1. European Center for Disease Prevention and Control. COVID-19 situation update worldwide. Available at: www.ecdc.europa.eu/en/geographical-distribution-2019-ncov-cases. Accessed 23 February.

2. Wolff D, Nee S, Hickey NS, Marschollek M. Risk factors for Covid-19 severity and fatality: a structured literature review. Infection 2021; 49(1): 15–28.

3. Pastor-Barriuso R, Perez-Gomez B, Hernan MA, et al. Infection fatality risk for SARS-CoV-2 in community dwelling population of Spain: nationwide seroepidemiological study. B.M.J. 2020; 371: m4509.

4. Meng Y, Wu P, Lu W, et al. Sex-specific clinical characteristics and prognosis of coronavirus disease-19 infection in Wuhan, China: A retrospective study of 168 severe patients. PLoS Pathog 2020; 16(4): e1008520.

5. Suleyman G, Fadel RA, Malette KM, et al. Clinical Characteristics and Morbidity Associated With Coronavirus Disease 2019 in a Series of Patients in Metropolitan Detroit. JAMA Netw Open 2020; 3(6): e2012270.

6. Chen T, Wu D, Chen H, et al. Clinical characteristics of 113 deceased patients with coronavirus disease 2019: retrospective study. B.M.J. 2020; 368: m1091.

7. Gupta S, Hayek SS, Wang W, et al. Factors Associated With Death in Critically Ill Patients With Coronavirus Disease 2019 in the US JAMA Intern Med 2020.

8. Takahashi T, Ellingson MK, Wong P, et al. Sex differences in immune responses that underlie COVID-19 disease outcomes. Nature 2020; 588(7837): 315–20.

9. Janssen NAF, Grondman I, de Nooijer AH, et al. Dysregulated innate and adaptive immune responses discriminate disease severity in COVID-19. J Infect Dis 2021.

10. Qin C, Zhou L, Hu Z, et al. Dysregulation of Immune Response in Patients With Coronavirus 2019 (COVID-19) in Wuhan, China. Clin Infect Dis 2020; 71(15): 762–8.

11. Wilk AJ, Rustagi A, Zhao NQ, et al. A single-cell atlas of the peripheral immune response in patients with severe COVID-19. Nat Med 2020; 26(7): 1070–6.

12. Yao C, Bora SA, Parimon T, et al. Cell type-specific immune dysregulation in severely ill COVID-19 patients. medRxiv 2020.

13. Mann ER, Menon M, Knight SB, et al. Longitudinal immune profiling reveals key myeloid signatures associated with COVID-19. Sci Immunol 2020; 5(51).

14. McClymont H, Hu W. Weather Variability and COVID-19 Transmission: A Review of Recent Research. Int J Environ Res Public Health 2021; 18(2).

15. Rayan RA Seasonal variation and COVID-19 infection pattern: A gap from evidence to reality. Curr Opin Environ Sci Health 2021; 20: 100238.

16. Aguirre-Gamboa R, Joosten I, Urbano P.C.M., et al. Differential Effects of Environmental and Genetic Factors on T and B Cell Immune Traits. Cell Rep 2016; 17(9): 2474–87.

17. Koeken VA, de Bree L.C.J., Mourits VP, et al. BCG vaccination in humans inhibits systemic inflammation in a sex-dependent manner. J Clin Invest 2020; 130(10): 5591–602.

18. Scully EP, Haverfield J, Ursin RL, Tannenbaum C, Klein SL. Considering how biological sex impacts immune responses and COVID-19 outcomes. Nat Rev Immunol 2020; 20(7): 442–7.

19. Chen Y, Klein SL, Garibaldi BT, et al. Aging in COVID-19: Vulnerability, immunity and intervention. Ageing Res Rev 2021; 65: 101205.

20. Huang W, Berube J, McNamara M, et al. Lymphocyte Subset Counts in COVID-19 Patients: A Meta-Analysis. Cytometry A 2020; 97(8): 772–6.

21. Chen G, Wu D, Guo W, et al. Clinical and immunological features of severe and moderate coronavirus disease 2019. J Clin Invest 2020; 130(5): 2620–9.

22. Akkaya M, Kwak K, Pierce SK. B cell memory: building two walls of protection against pathogens. Nat Rev Immunol 2020; 20(4): 229–38.

23. Bulut O, Kilic G, Dominguez-Andres J, Netea MG. Overcoming immune dysfunction in the elderly: trained immunity as a novel approach. Int Immunol 2020; 32(12): 741–53.

24. Sette A, Crotty S. Adaptive immunity to SARS-CoV-2 and COVID-19. Cell 2021; 184(4): 861–80.

25. Del Valle DM, Kim-Schulze S, Huang HH, et al. An inflammatory cytokine signature predicts COVID-19 severity and survival. Nat Med 2020; 26(10): 1636–43.

26. Komarowska I, Coe D, Wang G, et al. Hepatocyte Growth Factor Receptor c-Met Instructs T Cell Cardiotropism and Promotes T Cell Migration to the Heart via Autocrine Chemokine Release. Immunity 2015; 42(6): 1087–99.

27. Yang Y, Shen C, Li J, et al. Plasma IP-10 and MCP-3 levels are highly associated with disease severity and predict the progression of COVID-19. J Allergy Clin Immunol 2020; 146(1): 119–27 e4.

28. Schulte-Schrepping J, Reusch N, Paclik D, et al. Severe COVID-19 Is Marked by a Dysregulated Myeloid Cell Compartment. Cell 2020; 182(6): 1419–40 e23.

29. Blackburn SD, Wherry EJ. IL-10, T cell exhaustion and viral persistence. Trends Microbiol 2007; 15(4): 143–6.

30. Lu L, Zhang H, Dauphars DJ, He YW. A Potential Role of Interleukin 10 in COVID-19 Pathogenesis. Trends Immunol 2021; 42(1): 3–5.

31. Zhao Y, Qin L, Zhang P, et al. Longitudinal COVID-19 profiling associates IL-1RA and IL-10 with disease severity and RANTES with mild disease. JCI Insight 2020; 5(13).

32. Giamarellos-Bourboulis EJ, Netea MG, Rovina N, et al. Complex Immune Dysregulation in COVID-19 Patients with Severe Respiratory Failure. Cell Host Microbe 2020; 27(6): 992–1000 e3.

33. Bord S, Ireland DC, Beavan SR, Compston JE. The effects of estrogen on osteoprotegerin, RANKL, and estrogen receptor expression in human osteoblasts. Bone 2003; 32(2): 136–41.

34. Ono T, Hayashi M, Sasaki F, Nakashima T. RANKL biology: bone metabolism, the immune system, and beyond. Inflamm Regen 2020; 40: 2.

35. Lofgren E, Fefferman NH, Naumov YN, Gorski J, Naumova EN. Influenza seasonality: underlying causes and modeling theories. J Virol 2007; 81(11): 5429–36.

36. van der Made CI, Simons A, Schuurs-Hoeijmakers J, et al. Presence of Genetic Variants Among Young Men With Severe COVID-19. JAMA 2020.

37. Tang Y, Liu J, Zhang D, Xu Z, Ji J, Wen C. Cytokine Storm in COVID-19: The Current Evidence and Treatment Strategies. Front Immunol 2020; 11: 1708.

38. Gram AM, Frenkel J, Ressing ME. Inflammasomes and viruses: cellular defence versus viral offence. J Gen Virol 2012; 93(Pt 10): 2063–75.

39. Liu Y, Li S, Zhang G, et al. Genetic variants in IL1A and IL1B contribute to the susceptibility to 2009 pandemic H1N1 influenza A virus. BMC Immunol 2013; 14: 37.

40. Dinarello CA. Overview of the IL-1 family in innate inflammation and acquired immunity. Immunol Rev 2018; 281(1): 8–27.

